# New pathogenic variants and insights into pathogenic mechanisms in GRK1-related Oguchi disease

**DOI:** 10.1101/2020.02.20.936880

**Authors:** James A. Poulter, Molly S. C. Gravett, Rachel L. Taylor, Kaoru Fujinami, Julie De Zaeytijd, James Bellingham, Atta Ur Rehman, Takaaki Hayashi, Mineo Kondo, Abdur Rehman, Muhammad Ansar, Dan Donnelly, Carmel Toomes, Manir Ali, UK Inherited Retinal Disease Consortium, Genomics England Research Consortium, Elfride De Baere, Bart P. Leroy, Nigel P. Davies, Robert H. Henderson, Andrew R. Webster, Carlo Rivolta, Omar A. Mahroo, Gavin Arno, Graeme C. Black, Martin McKibbin, Sarah A. Harris, Kamron N. Khan, Chris F. Inglehearn

## Abstract

**Purpose:** Biallelic mutations in G-Protein coupled receptor kinase 1 (GRK1) cause Oguchi disease, a rare subtype of congenital stationary night blindness (CSNB). The purpose of this study was to identify pathogenic GRK1 variants and use in-depth bioinformatic analyses to evaluate how their impact on protein structure could lead to pathogenicity.

**Methods:** Patients’ genomic DNA was sequenced by whole genome, whole exome or focused exome sequencing. Pathogenic variants, published and novel, were compared to nondisease associated missense variants. The impact of *GRK1* missense variants at the protein level were then predicted using a series of computational tools.

**Results:** We identified eleven previously unpublished cases with biallelic pathogenic GRK1 variants, including seven novel variants, and reviewed all *GRK1* pathogenic variants. Further structure-based scoring revealed a hotspot for missense variants in the kinase domain. Additionally, to aid future clinical interpretation, we identified the bioinformatics tools best able to differentiate pathogenic from non-pathogenic variants.

**Conclusion:** We identified new *GRK1* pathogenic variants in Oguchi disease patients and investigated how disease-causing variants may impede protein function, giving new insights into the mechanisms of pathogenicity. All pathogenic GRK1 variants described to date have been collated into a Leiden Open Variation Database (http://dna2.leeds.ac.uk/GRK1_LOVD/genes/GRK1).

## Introduction

The first member of the G protein-coupled receptor kinase (GRK) family was discovered when enzymatic activity was observed in rod membranes that phosphorylated rhodopsin in a light-dependent manner (Kuhn & Dreyer, 1972). The enzyme, now known as GRK1 (MIM 180381), was found to be essential for quenching and recycling light-activated rhodopsin. To be recycled, light-activated rhodopsin is phosphorylated multiple times by GRK1, followed by binding of Arrestin-1 to activated-phosphorylated rhodopsin to block further transducin activation by steric exclusion (Krupnick, Gurevich, & Benovic, 1997; Wilden, Hall, & Kuhn, 1986). Failure of this process results in a build-up of activated rhodopsin and a lack of light-sensitive rhodopsin available for detection of light. Importantly, a build-up of activated rhodopsin appears to be well tolerated by photoreceptors, with no obvious cell death or structural consequences in the retina, meaning any future therapies leading to the deactivation of rhodopsin could restore visual acuity. Activated cone opsin is likely to be more reliant on GRK7 than GRK1 phosphorylation, as although both enzymes are expressed in cones, loss of GRK1 function impacts minimally on human photopic vision.

GRK1 is a serine/threonine protein kinase with a central catalytic AGC protein kinase (PK) domain that sits within a regulator of G protein signalling homology (RH) domain (Fig. 1). The PK domain binds ATP and the polypeptide substrate which is subsequently phosphorylated (Arencibia, Pastor-Flores, Bauer, Schulze, & Biondi, 2013). The RH domain is thought to have a key role in receptor binding, with loss of this domain preventing binding and phosphorylation of rhodopsin (He et al., 2017). While the N-terminus encodes a short alpha-helical domain, the C-terminus encodes a lipid-binding region, which is crucial for prenylation-dependent docking of GRK1 into the outer segment membranes of rod photoreceptors (Komolov & Benovic, 2018).

**Figure 1.**
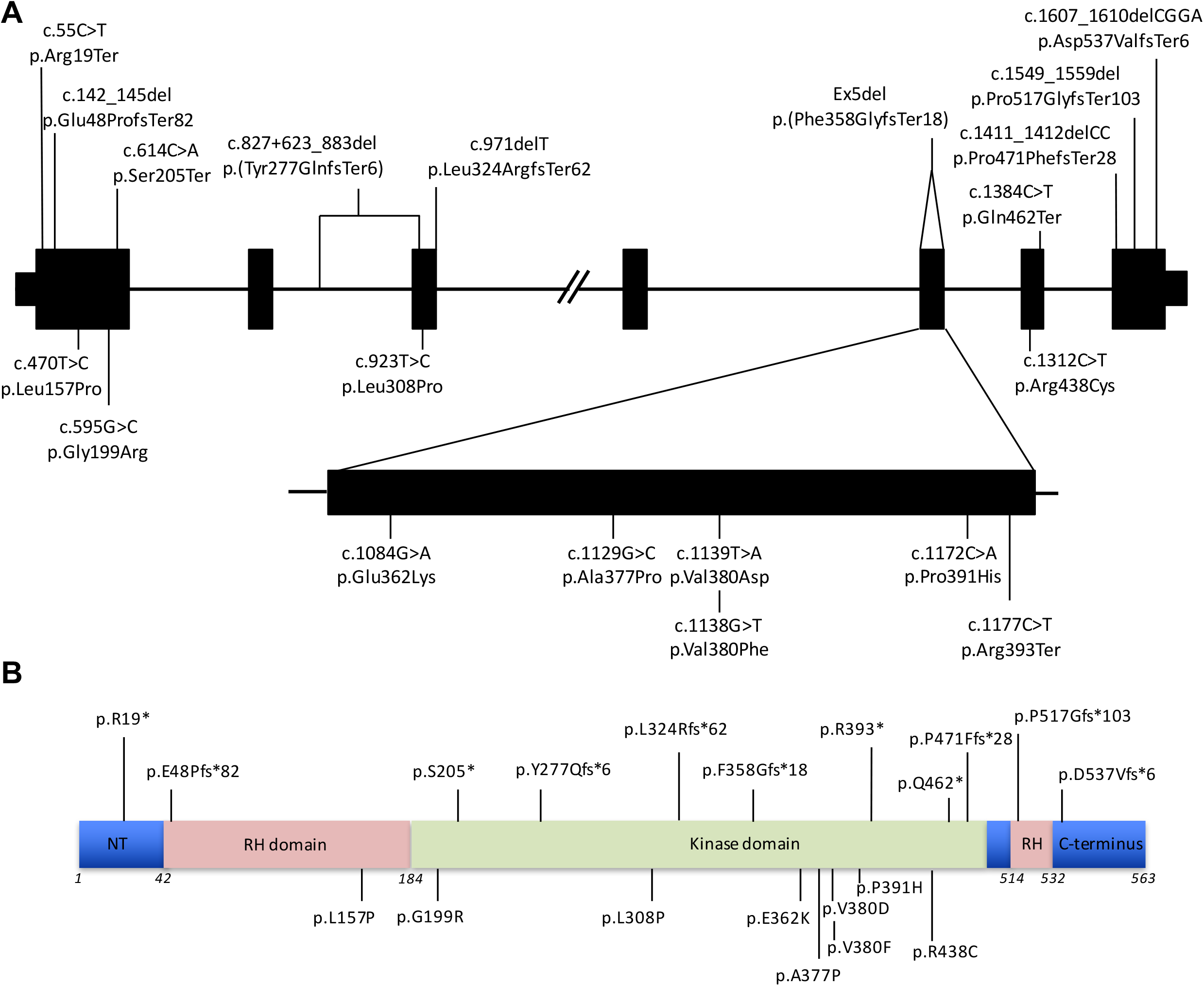
Distribution of pathogenic variants within GRK1. All variants shown are annotated according to human genome GRCh37/hg19, using GRK1 gene and protein accession numbers NM_002929 and NP_002920 respectively. Loss of function variants are shown above the gene/protein and missense variants are given below. (A) Genomic organisation of *GRK1* showing the location of all novel and published Oguchi disease causing variants. (B) Domain structure of GRK1 showing the location of all GRK1 variants. The start and end amino-acid positions of each domain are based on Lodowski *et al.* (2006)(Lodowski, Tesmer, Benovic, & Tesmer, 2006). NT = N-Terminus, RH = regulator of G-protein signalling homology.

Oguchi disease (MIM 613411), a rare form of congenital stationary night blindness (CSNB), results from biallelic variants in either *SAG* (encoding Arrestin-1) or *GRK1. SAG*-mediated disease is most prevalent in Japanese patients, whilst pathogenic variants in *GRK1* are more common in South East Asians. Typically, Oguchi disease is characterised clinically by the Mizuo-Nakamura phenomenon – the presence of a golden-yellow colouration to the retina that disappears in the dark-adapted state and reappears shortly after light exposure (Miyake, Horiguchi, Suzuki, Kondo, & Tanikawa, 1996). Electrophysiologically there is normal cone function, delayed rod dark adaptation and marked rod desensitisation to a bright flash. Most reported cases have this distinctive phenotype and no significant variation in disease expression has been reported, with the exception of a few rare cases (Hayashi, Gekka, Takeuchi, Goto-Omoto, & Kitahara, 2007; Nishiguchi et al., 2019). Distinguishing between Oguchi disease and other forms of CSNB is important as they may have different prognoses – some patients with Oguchi disease report very slowly progressive visual dysfunction, whereas those with classical CSNB do not. Minimally progressive retinal degeneration has been observed in *SAG*-associated Oguchi disease, and for one patient, this resulted in their disease being re-classified as retinitis pigmentosa 26 years after being diagnosed with a stationary rod dysfunction syndrome (Oguchi disease)(Nishiguchi et al., 2019). As most of our knowledge of Oguchi disease stems from our understanding of loss of SAG function further research is required in order to better understand *GRK1*-related Oguchi disease.

To date, thirteen pathogenic variants in *GRK1* have been implicated in Oguchi disease, eight of which are predicted to be null variants (Azam et al., 2009; Cideciyan et al., 1998; Godara et al., 2012; Hayashi et al., 2007; Jespersgaard et al., 2019; Li et al., 2017; Mucciolo et al., 2018; Oishi et al., 2007; Skorczyk-Werner, Kociecki, Wawrocka, Wicher, & Krawczyniski, 2015; Teke, Citirik, Kabacam, Demircan, & Alikasifoglu, 2016; Yamamoto, Sippel, Berson, & Dryja, 1997; Zhang et al., 2005). The molecular mechanisms by which these contribute to disease are poorly understood. Here we present seven further pathogenic variants in *GRK1*, and review new and known variants causing Oguchi disease. Using *in-silico* techniques we compare known disease-associated, with likely non-pathogenic, *GRK1* missense variants to identify features that could be contributing to the phenotype. Together this will help define disease mechanisms and assist in predicting likelihood of pathogenicity for variants identified in *GRK1*.

## Methods

### Editorial Policies and Ethical Considerations

Ethical approval for this study was obtained from the Yorkshire and The Humber – Leeds East Research Ethics Committee (REC reference: 17/YH/0032). The study adhered to the tenets of the Declaration of Helsinki. Blood samples were taken with informed consent from each participant, or with parental informed consent on behalf of children.

### Clinical assessment

Study participants were ascertained from St James’ University Hospital, Leeds, England; Manchester Centre for Genomic Medicine, Manchester, England; Moorfields Eye Hospital, London, England; Department of Ophthalmology, Ghent University Hospital, Ghent, Belgium; The Jikei University School of Medicine, Tokyo, Japan; Mie University Graduate School of Medicine, Mie, Japan; and National Institute of Sensory Organs, National Hospital Organization Tokyo Medical Centre, Tokyo Japan; Individuals were recruited following clinical examination by an experienced ophthalmologist, with the exception of Families 10 and 11 who were ascertained by local doctors in Pakistan.

### Sequencing

Exome and clinical exome sequencing was performed on 3μg of genomic DNA using the Agilent SureSelectXT Human All Exon (V6) and Focussed Exome respectively (Agilent Technologies, CA, USA), according to the manufacturer’s instructions. Captured libraries were pooled and sequenced on an Illumina HiSeq3000 sequencer (Illumina, CA, USA). The resulting fastq files were aligned to the Human genome assembly GRCh37 using Burrows-Wheeler Aligner (BWA) and reads processed using samtools, Picard and the Genome Analysis Toolkit. Variants within the captured regions were called and annotated using variant effect predictor (NCBI) and further filtered for those within genes known to cause retinal disease. This analysis pipeline is described in more detail elsewhere (Panagiotou et al., 2017). Copy number variation (CNV) analysis was performed using ExomeDepth, comparing patient BAM files with BAM files from unrelated individuals in the same sequencing run. All novel variants were confirmed by Sanger sequencing using standard methods.

The affected individual and unaffected parents of Family 1 underwent Genome sequencing as part of the 100,000 Genome Project, as previously detailed by Taylor *et al* (Taylor et al., 2017).

### Identifying non-pathogenic variants

To identify protein-level characteristics influencing a disease phenotype we compared the 9 pathogenic missense variants with a group of 5 assumed non-pathogenic missense variants. For the non-pathogenic group, we identified all homozygous *GRK1* missense variants in gnomAD and only included those with a MAF >0.00035, the level above which variants are calculated to be too common to cause Oguchi disease (Whiffin et al., 2017). The five missense variants in gnomAD with frequencies above this level, p.Leu54Phe, p.Thr97Met, p,Ile241Thr, p.Glu464Gln and p.Ser536Leu, also all occur at least once as homozygotes in gnomAD. These are therefore assumed to be non-disease associated polymorphisms (further details in Supplementary Table 1), which for the purposes of this paper we will call non-pathogenic variants.

### Structural analysis

A homology model of human wild-type GRK1 based on the bovine structure (PDB: 4PNI) was produced using SwissModel (Waterhouse et al., 2018). The locations of the RH and PK domains, ATP binding site, polypeptide substrate binding site and activation loop were identified using InterPro (Mitchell et al., 2019). The structure was visualised, and regions of interest labelled using PyMOL (Schrodinger, 2015). Residue depth and connectivity within the structure was determined using Site Directed Mutator (SDM) (Pandurangan, Ochoa-Montano, Ascher, & Blundell, 2017). To further explain variant impact, changes in residue physical characteristics; molecular weight (Mw), isoelectric point (pI) and hydrophobicity (Laskowski, Stephenson, Sillitoe, Orengo, & Thornton, 2020) were reviewed.

### Scoring missense variants

A number of bioinformatics tools were reviewed to identify which approach best classified the variants. Scoring methods included: SDM (Pandurangan et al., 2017), VarSite (Laskowski et al., 2020), SIFT (Sim et al., 2012), Polyphen-2 (Adzhubei et al., 2010), CADD (Kircher et al., 2014), Rhapsody (Ponzoni, Oltvai, & Bahar, 2019) and Consurf (Ashkenazy et al., 2016). All statistical analyses were performed in GraphPad Prism 8.0.2 (for macOS, GraphPad Software, San Diego, California USA, www.graphpad.com) using Welch’s T-test.

## Resuslts

### Identification of novel *GRK1* pathogenic variants

Screening of inherited retinal disease patients by an international consortium of retinal genetics laboratories identified eleven cases with Oguchi disease harbouring eight different pathogenic variants in *GRK1*, of which seven are previously unreported (Table 1). All cases are of South Asian ethnicity except for Family 2 who are East Asian (Japanese) and Family 5 who are Eastern European (Polish). All variants were confirmed and segregated in available family members. Each patient had a history of non/minimally-progressive night blindness, and retinal examination revealed the Mizuo-Nakamura phenomenon, consistent with a clinical diagnosis of Oguchi disease.

**Table 1.** Summary of all novel and published mutations in GRK1.

| DNA variant | Exon | Protein variant | rs number | gnomAD Frequency | Domain | PolyPhen-2 | CADD (v1.3) | References |
| --- | --- | --- | --- | --- | --- | --- | --- | --- |
| c.55C>T | 1 | p.Arg19Ter | rs370713047 | 17/239,700<br>(1 homozygous) | N-Terminus | N/A | 37.0 | Li et al (2017) |
| c.142_145del | 1 | p.Glu48ProfsTer82 | rs748680704 | 6/248,384 | RH domain | N/A | 28.5 | This study (**F1, F8**) |
| c.470T>C | 1 | p.Leu157Pro | N/A | 0 | RH domain | 1.00<br>(Prob. D) | 26.0 | Mucciolo et al (2018) |
| c.595G>C | 1 | p.Gly199Arg | N/A | 0 | Kinase | 1.00<br>(Prob. D) | 24.7 | This study (**F2**) |
| c.614C>A | 1 | p.Ser205Ter | N/A | 0 | Kinase | N/A | 36 | Azam et al (2009) |
| c.827+623_883del | 3 | p.(Tyr277GlnfsTer6) | N/A | 0 | Kinase | N/A | 27.8 | Zhang et al (2005) |
| c.923T>C | 3 | p.Leu308Pro | N/A | 0 | Kinase | 0.996<br>(Prob. D) | 29.2 | Teke et al. (2016) |
| c.971del | 3 | p.Leu324ArgfsTer62 | N/A | 0 | Kinase | N/A | 35 | Oishi et al (2007) |
| Ex5 del | 5 | p.(Phe358GlyfsTer18) | N/A | 0 | Kinase | N/A | 25.1<sup>a</sup> | Yamamoto et al. (1997);<br>Cideciyan et al. (1998) |
| c.1084G>A | 5 | p.Glu362Lys | N/A | 0 | Kinase | 1.00<br>(Prob. D) | 34 | This study (**F3**) |
| c.1129G>C* | 5 | p.Ala377Pro | N/A | 0 | Kinase | 0.991<br>(Prob. D) | 25 | Godara et al (2012)<sup>1</sup> |
| c.1138G>T | 5 | p.Val380Phe | N/A | 2/173,226 | Kinase | 0.998<br>(Prob. D) | 32 | This study (**F4**) |
| c.1139T>A* | 5 | p.Val380Asp | rs777094000 | 5/173,124 | Kinase | 0.999<br>(Prob. D) | 34 | Yamamoto et al. (1997)<sup>2</sup>;<br>Godara et al. (2012)<sup>1</sup> |
| c.1172C>A | 5 | p.Pro391His | rs570621429 | 1/141,802 | Kinase | 1.00<br>(Prob. D) | 34 | Hayashi et al (2007) |
| c.1177C>T | 5 | p.Arg393Ter | rs137877289 | 6/141,764 | Kinase | N/A | 46 | This study (**F5**)<sup>3</sup> |
| c.1312C>T | 6 | p.Arg438Cys | rs765070399 | 2/141,922 | Kinase | 1.00<br>(Prob. D) | 34 | This study (**F9**) |
| c.1384C>T | 6 | p.Gln462Ter | N/A | 0 | Kinase | N/A | 42 | Jespersgaard et al (2019) |
| c.1411_1412del | 7 | p.Pro471PhefsTer28 | N/A | 0 | Kinase | N/A | 35 | Oishi et al (2007) |
| c.1549_1559del | 7 | p.Pro517GlyfsTer130 | N/A | 0 | RH domain | N/A | 35 | This study (**F6**) |

**Table 1 – Summary of all novel and published mutations in GRK1.**
|  |  |  |  |  |  |  |  |  |
| --- | --- | --- | --- | --- | --- | --- | --- | --- |
| c.1607_1610del | 7 | p.Asp537ValfsTer7 | rs756235051 | 89/172,958 | C-Terminus | N/A | 26.7 | Yamamoto et al (1997)<sup>2</sup>; Skorczyk-Werner et al (2015); This study<br>**(F5<sup>3</sup>, F7, F10, F11)** |

All mutations are shown relative to Human genome reference GRCh37, and GRK1 references NM\_002929 and NP\_002920. The precise start and end locations for each domain in GRK1 were obtained from Lodowski et al. (2006). All variants identified are all homozygous except for <sup>1,2,3</sup>, which are compound heterozygous, \* Predicted – DNA mutation not provided in publication, Prob. D = Probably Damaging as scored by PolyPhen2. <sup>a</sup> Based on deletion of whole of exon 5 and surrounding splice sites as exact coordinates are unknown.

A literature search revealed a further thirteen published Oguchi disease causing *GRK1* variants from twelve studies (Azam et al., 2009; Cideciyan et al., 1998; Godara et al., 2012; Hayashi et al., 2007; Jespersgaard et al., 2019; Li et al., 2017; Mucciolo et al., 2018; Oishi et al., 2007; Skorczyk-Werner et al., 2015; Teke et al., 2016; Yamamoto et al., 1997; Zhang et al., 2005). The nomenclature for these variants was updated based on the current HGVS nomenclature guidelines and the GRCh37 version of the Human Genome, and together with the new variants reported here, these have been included in a Leiden Open Variation Database (LOVD) of *GRK1* variants (http://dna2.leeds.ac.uk/GRK1_LOVD/genes/GRK1).

### Pathogenic variant distribution and domain structure

The twenty Oguchi disease causing *GRK1* variants identified to date include missense (n=9), frameshift (n=6) and nonsense (n=4) variants (Table 1) as well as a large deletion encompassing exon 5 (Fig. 1A). Each variant has been identified in only a single family with the exception of p.Val380Asp (2 families), p.Glu48ProfsTer82 (2 families), the deletion of exon 5 (p.(Phe358GlyfsTer18)) (3 families) and the C-terminal frameshift variant p.(Asp537ValfsTer6) (6 families), with the latter variant having the highest allele frequency in gnomAD (89/172,958). Of the 89 alleles in gnomAD, 42 are in European (non-Finnish) and 35 in South Asian populations, which is consistent with this mutation only being identified in cases from Eastern Europe and South Asia.

Fifteen of the variants lie within the protein kinase domain, three within the RH domain and one at each of the N- and C-termini (Fig. 1B). While the frameshift and nonsense variants are present throughout the protein, all except one of the missense variants (n=8 out of 9) are present within the protein kinase domain, and five cluster within a 30 amino-acid region between Glu362 and Pro391. These are likely to impair phosphorylation of light-activated rhodopsin.

Three variants located after this cluster introduce frameshifts in the final exon, giving rise to transcripts which would not be expected to undergo nonsense-mediated decay (Thermann et al., 1998). The extreme C-terminus of GRK1, which these three variants would disrupt, is prenylated to facilitate anchoring of GRK1 in the membranes of the rod photoreceptor outer segments (Pitcher, Freedman, & Lefkowitz, 1998), which maximizes the likelihood of interaction with activated rhodopsin. Lack of this domain is therefore likely to reduce the level of anchored GRK1, which will in turn reduce the quantity of rhodopsin being bound and phosphorylated.

### Pathogenic missense variants affect key regions in *GRK1* active site

To further understand the mechanism by which the missense variants affected GRK1 function, we performed structure-based scoring on the SDM server using our homology model (Fig. 2A-C). Although SDM’s primary function is predicting thermal stability, the most interesting results came from the scoring of wildtype (WT) residue depth and occluded surface packing (OSP), with both scores showing a significant difference between pathogenic and non-pathogenic variants (p<0.01 and p<0.05 respectively). This demonstrated that pathogenic variants were located deeper within the 3D structure of *GRK1* than non-pathogenic variants (residue depth = 3.5-11.8 and 3-6.2 respectively) (Fig. 2A, Supplementary Table 2) and were more connected/densely packed (OSP = 0.3-0.57 and 0.04-0.48 respectively) (Fig. 2B, Supplementary Table 2). Exceptions to these rules were seen in p.Gly199Arg (pathogenic) which is located at the surface of the protein and p.Ile241Thr (non-pathogenic) which is located deeper in GRK1. Gly199 is positioned within the conserved glycine rich loop of the PK domain and is part of the ATP binding site (Fig. 3). Mutation to a bulky positively charged arginine (Supplementary Table 2) is likely to disrupt both the shape and charge of the ATP binding pocket (Fig. 3). Although Ile241 is buried in the binding pocket of the PK domain (Fig. 3), there is little change in size and charge (Supplementary Table 2) which could explain why this variant is non-pathogenic.

**Figure 2.**
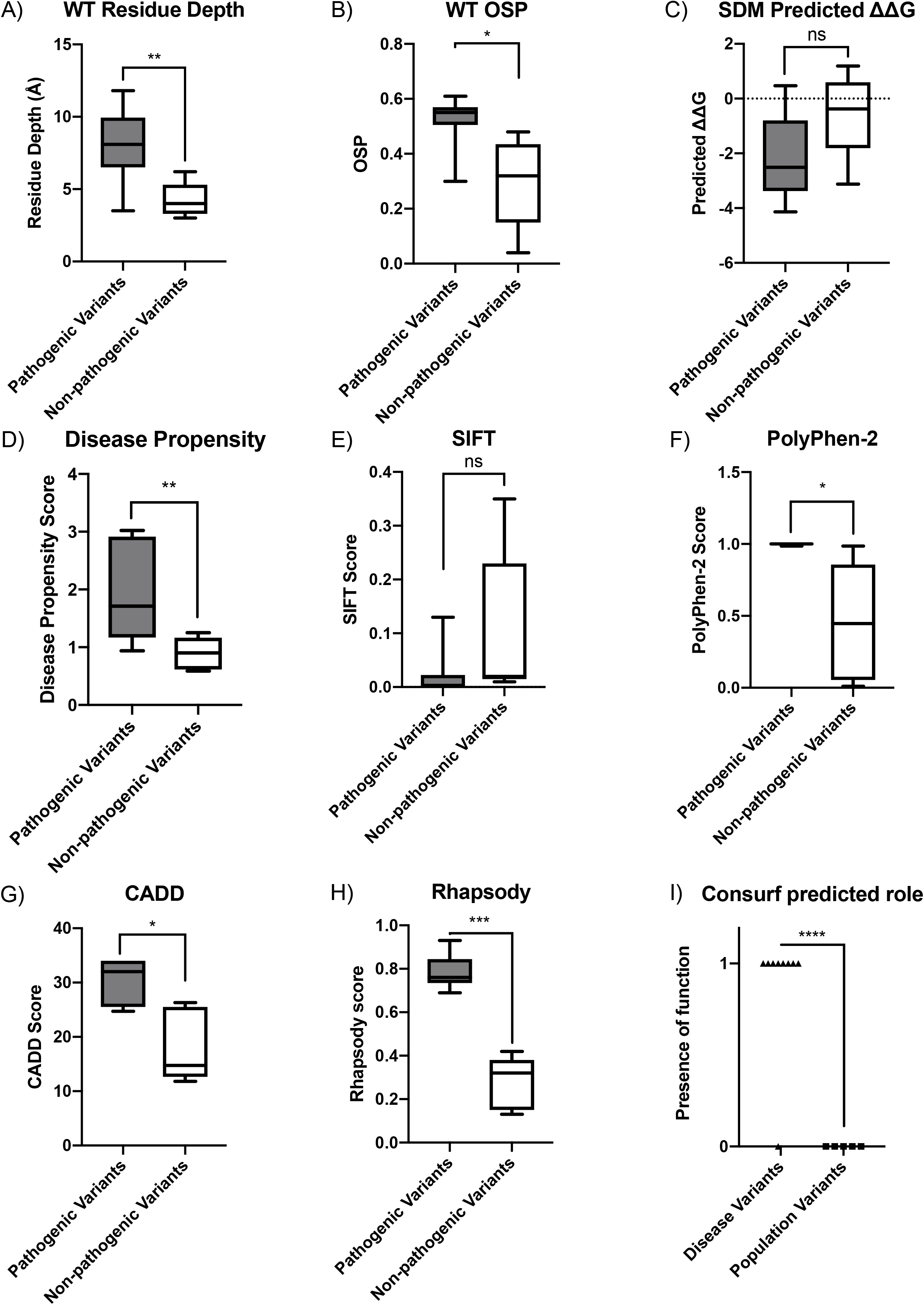
Bioinformatic prediction scores of pathogenic and non-pathogenic GRK1 variants. Comparison of predicted pathogenicity scores for all novel or published pathogenic missense GRK1 variants, with likely non-pathogenic missense variants identified in gnomAD. The whiskers in the box plots show the minimum and maximum values. Statistical significance was calculated using Welch’s T-test in GraphPad prism 8.0.2. ns = p ≥0.05, * = p<0.05, ** = p<0.01, *** = p<0.001. (A) Wildtype (WT) residue depth from Site Directed Mutator (SDM), (B) WT occluded surface packing (OSP) from SDM, (C) Thermostability score SDM, (D) Disease propensity scores as predicted in VarSite, (E) SIFT scores, (F) PolyPhen-2 scores, (G) CADD (v1.3) scores, (H) Rhapsody scores and (I) Consurf predicted roles (1 = structural/functional role predicted, 0 = no role predicted).

**Figure 3.**
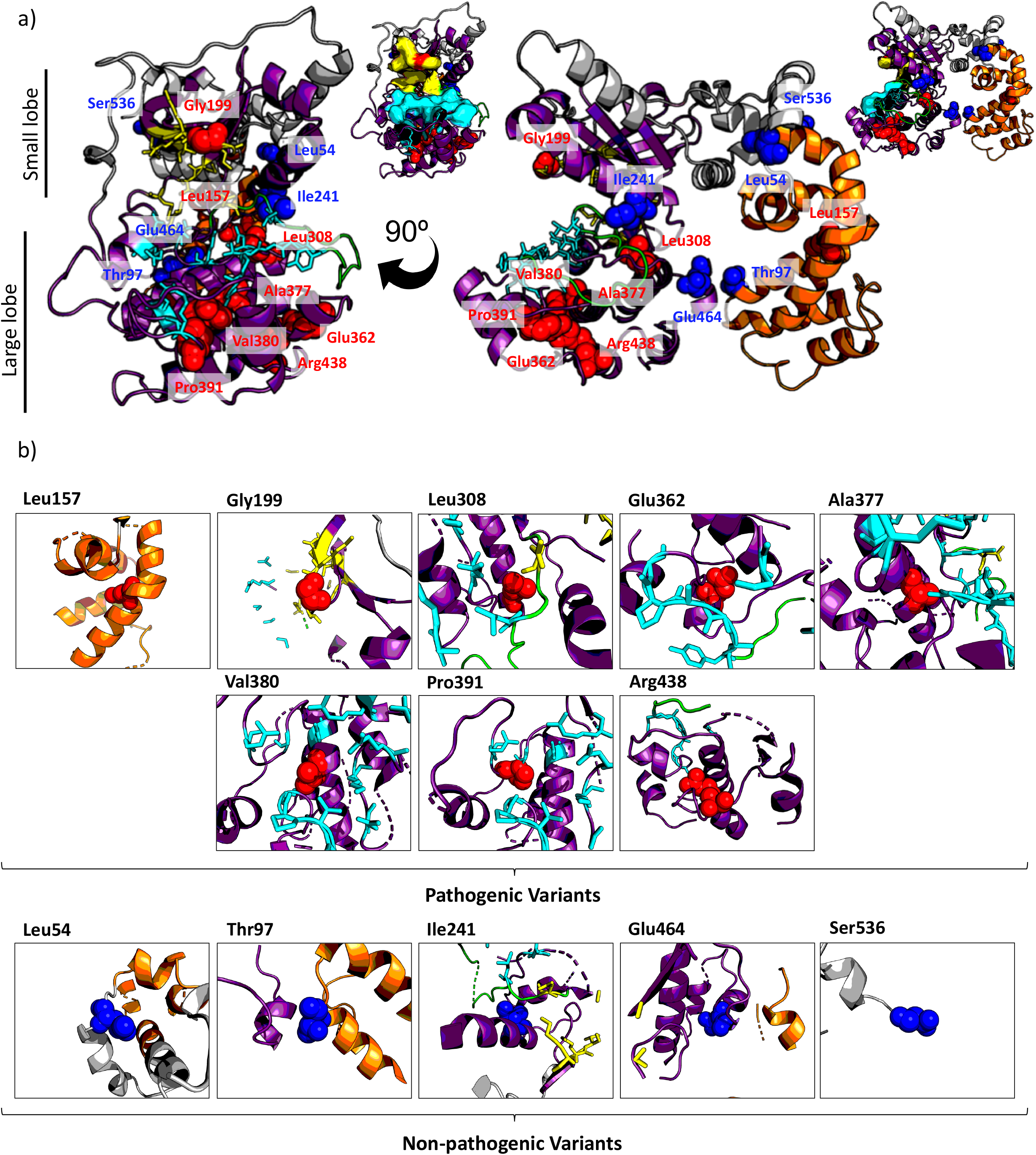
Pathogenic and non-pathogenic variants location in homology model of GRK1. Homology model of human GRK1 based upon bovine GRK1 structure (PDB: 4PNI). Wildtype residues of pathogenic variants are labelled as red spheres (Leu157, Gly199, Leu308, Glu362, Ala377, Val380, Pro391 and Arg438) and non-pathogenic as blue spheres (Leu54, Thr97, Ile241, Glu464 and Ser536). The protein kinase domain is coloured in purple, RH domain in orange, activation loop in green, ATP binding residues are represented as yellow sticks and polypeptide substrate binding residues as cyan sticks. (a) A visualisation of variant location in the overall structure. The smaller representations to the right, with the ATP and polypeptide binding surface displayed demonstrate how intramolecular residue changes may deform the binding site surface. (b) A focus on residue location and local interactions, displaying only features within 15Å of the residue of interest.

Further investigation of mutation location in the protein structure demonstrated pathogenic variants were located proximal to key sites for protein function, including the substrate and ATP binding domains, while non-pathogenic variants were not (Fig. 3A). Pathogenic variants were predicted to be likely to disrupt the shape and charge of the active site either directly as in the case of p.Gly199Arg, or allosterically through introduction of proline kinks into alpha helices (e.g. p.Leu308Pro & p.Ala377Pro), intramolecular charged groups (classically exposed) (e.g. p.Val380Asp & p.Pro391His), and bulky side groups within densely packed regions (e.g. p.Val380Phe) close to key residues (Fig. 3B, Supplementary Table 2). In the case of p.Arg438Cys and p.Glu362Lys, the equivalent arginine and glutamic acid residues are conserved and form a salt bridge in the bovine protein (PDB: 4PNI). It is therefore likely that substitution of Glu362 and Arg438 changes the charge of these residues in the human protein and is disrupting this salt bridge interaction (Supplementary Figure 2). Meanwhile non-pathogenic variants were often subtler changes, and not local to key residues (Fig. 3B, Supplementary Table 2). We hypothesise pathogenic variants within the protein kinase domain are inhibiting protein function through disruption of the shape and dynamics of this key region, while non-pathogenic variants prove less disruptive to shape as they are not internal and/or have subtler changes in residue characteristics. This is supported by Consurf’s prediction that most of the pathogenic variants occur at loci important for protein structure or function (Fig. 2I, Supplementary Table 2).

Only one pathogenic missense variant, p.Leu157Pro, lies outside of the kinase domain, instead being present within the RH domain of GRK1, the domain primarily responsible for rhodopsin binding (He et al., 2017). The introduction of a proline residue within an α-helix is likely to disrupt the secondary structure and introduce a characteristic “kink” due to its rigidity and constrained phi angle. This could potentially impact on the ability of Rhodopsin to correctly bind GRK1 in order for phosphorylation to occur.

To determine which bioinformatic tools were most successful in differentiating between pathogenic and non-pathogenic variants, we scored missense variants using: VarSite, SDM, PolyPhen-2, SIFT, CADD, Rhapsody and Consurf. Predictions were compared between pathogenic and non-pathogenic variant sets, with significance assessed by Welch’s T-test (Fig 2, Table 1, Supplementary Table 2). VarSite disease propensity, WT OSP, WT residue depth, PolyPhen-2, CADD, Rhapsody and Consurf, were able to distinguish between pathogenic and non-pathogenic variants with statistical significance. Consurf’s structure/function prediction and Rhapsody, which both take the variant’s location in the protein 3D structure into consideration, were able to differentiate between pathogenic and non-pathogenic most successfully (p <0.001) (Fig. 2H&I, Supplementary Table 2).

Rhapsody has the additional feature of being able to predict the impact of substituting any residue in a protein with all of the 19 other amino acids *(in silico* site directed mutagenesis), making it extremely valuable diagnostically, as it has the potential to inform future work by identifying residue changes which are likely to be poorly tolerated. We performed protein-wide site directed mutagenesis and reviewed the top scoring variants (Rhapsody score ≥ 0.896). This analysis revealed that the top scoring variants were present at just 22 loci across the protein, thus identifying a number of key residues with the potential to cause Oguchi disease, if mutated. Of the 22 amino acid residues that Rhapsody predicts could harbour the most damaging variants, 20 are located in the protein kinase domain (Fig. 4).

**Figure 4.**
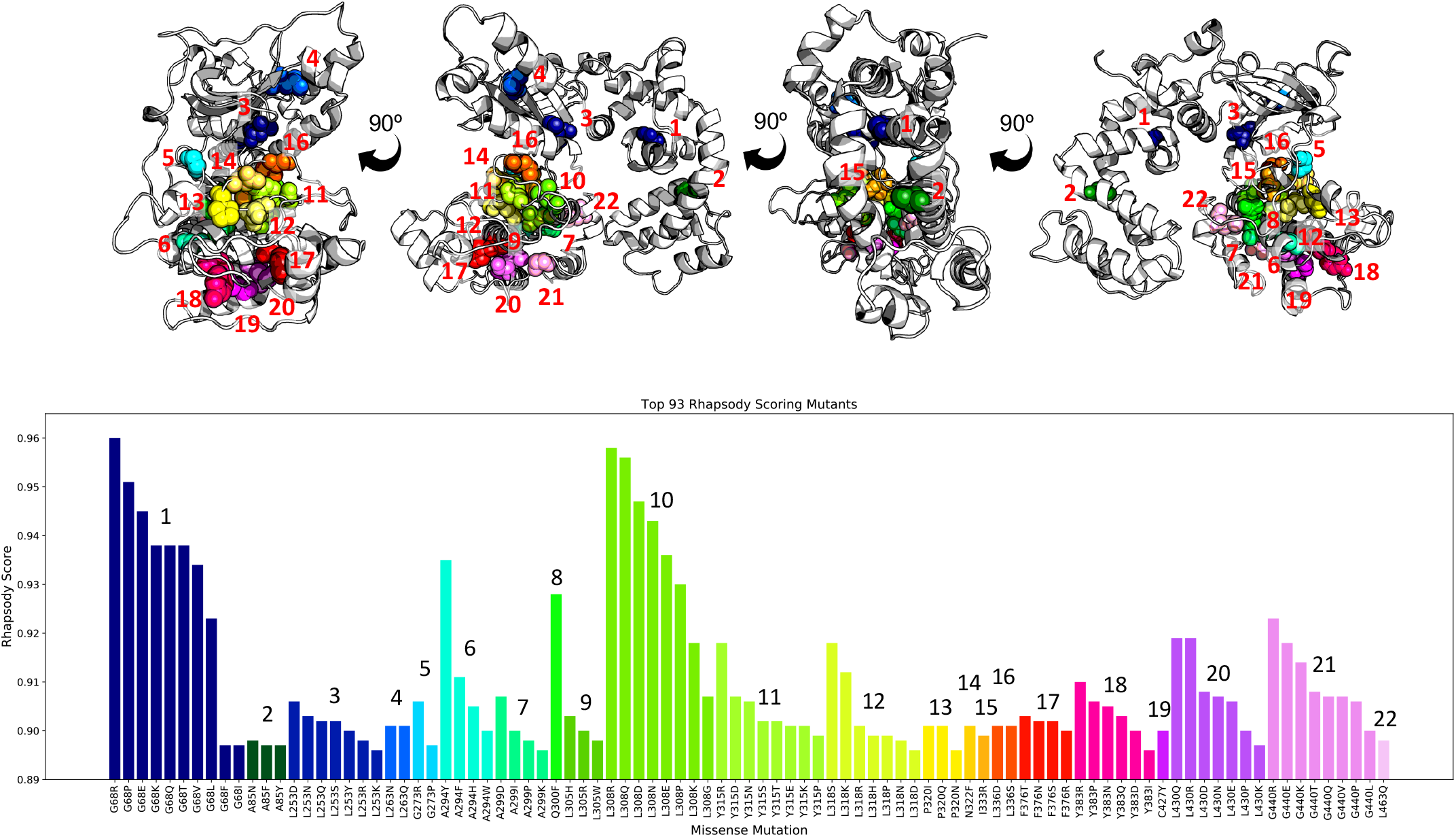
Rhapsody top scoring substitutions mapped onto the homology model of GRK1. Homology model of human GRK1 based upon bovine GRK1 structure (PDB: 4PNI). Wildtype residues of top-scoring ((≥0.896) rhapsody variants are labelled as spheres, coloured and numbered based on residue number. The colours and labels within the bar chart of top-scoring Rhapsody variants mimic this format. Locations with >4 variants in the top 93 scores include: G68, L253, L308, Y315, L318, Y383, L430 and G440.

## Discussion

Using a combined genetics and structural biology approach we have identified individuals with Oguchi disease due to biallelic *GRK1* variants and inferred the likely functional consequences of these and other published variants. We identified seven new pathogenic variants, taking the total number of *GRK1* variants associated with Oguchi disease to twenty. While eleven result in premature termination codons, nine result in single amino acid substitutions.

It is likely that transcripts encoding nonsense and frameshift variants, except those in the last exon, will be subject to nonsense mediated decay (NMD)(Thermann et al., 1998). This has not been confirmed in patient cells, but even if a truncated GRK1 lacking the residues encoded by exon 5 were produced, function has been shown to be significantly reduced, resulting in protein that was not able to phosphorylate Rhodopsin (Cideciyan et al., 1998). This implies that lack of functional GRK1 protein is one mechanism by which Oguchi disease can arise.

eight of the nine observed missense variants lie within the protein kinase domain, which is critical for GRK1 function. The kinase domain is the largest within GRK1, and variants in this region are likely to have a negative impact on phosphorylation of light-activated rhodopsin. Indeed, a previous study has shown the p.Val380Asp variant has no kinase activity for Rhodopsin compared to wild-type GRK1 (Khani, Nielsen, & Vogt, 1998). Five missense variants appear to be clustered within a 30 amino-acid region between Glu362 and Pro391, which may imply that amino-acid substitutions in this region are more likely to disrupt protein function than those located elsewhere. These variants were predicted to affect the structure of GRK1 and therefore kinase activity. As the structural analysis relies on a homology model of human GRK1, it is not always feasible to quantify atomic level changes (e.g. individual H-bond or salt bridge interactions within non-conserved residues). However, it is possible to draw general conclusions about the surrounding residue location and impact on secondary and tertiary structure. Therefore, loss of kinase activity through disruption of its shape and dynamics is a second mechanism by which Oguchi disease can occur.

Three variants have been identified in the last exon of *GRK1*, all of which result in frameshift transcripts that potentially escape nonsense mediated decay, leading to a GRK1 protein with a disrupted C-terminus. This region plays a crucial role in embedding GRK1 within the photoreceptor outer segment membrane; GRK1 localises directly to the outer segment membranes due to a short prenylation sequence within the C-terminus that interacts with the prenyl-binding protein, delta (PrBP/δ), which facilitates attachment to the membranes (Huang, Orban, Jastrzebska, Palczewski, & Tesmer, 2011; Roosing, Collin, den Hollander, Cremers, & Siemiatkowska, 2014). This cellular localisation is essential for GRK1 function as it brings active rhodopsin into contact with docked GRK1 protein through a twodimensional search rather than three-dimensional diffusion, which is essential for ultra-rapid signal termination (Sato, Chuprun, Schwartz, & Koch, 2015). Lack of this sequence is likely to cause failure of prenylation, leading to abolished binding of PrBP/δ and incorrect localisation of GRK1, a third mechanism by which disease can arise.

Our study therefore highlights three potential molecular mechanisms by which GRK1 variants lead to disease, each of which ultimately reduce or abolish the phosphorylation of light-activated rhodopsin. These are a) lack of GRK1 protein due to frame shift or nonsense variants in all exons but the last, resulting in NMD; b) missense variants leading to inability of GRK1 to phosphorylate rhodopsin due to a dysfunctional kinase domain or failure to bind rhodopsin, ATP or Mg2+; or c) a frameshift in the final exon, which would be expected to escape nonsense mediated decay and produce a protein retaining the ability to bind and phosphorylate light-activated rhodopsin, but which cannot localise to the outer segment membranes. We would therefore predict that, while variants in the first category lead to no protein being produced, those in the second produce proteins that correctly localise within the outer segments but have inhibited kinase activity, and those in the third produce a GRK1 protein likely to maintain kinase activity but which fails to localise and so rarely comes into contact with activated rhodopsin. Interestingly, the kinase activity of the p.(Asp537ValfsTer6) variant was found to be significantly reduced in transfected cos7 cells, suggesting c-terminal mutations may affect kinase activity as well as protein localisation. However, this lack of activity may be explained by difficulties in expressing the mutation containing GRK1 protein rather than on protein function (Khani et al., 1998).

Our comparison of GRK1 pathogenic variants to non-pathogenic variants identified from gnomAD facilitated an assessment of pathogenicity scores, which could inform future variant interpretation. We found that VarSite disease propensity, Polyphen2, CADD, Rhapsody and Consurf were able to significantly differentiate between pathogenic and non-pathogenic variants, as were biophysical scores of residue depth and occluded surface packing. Rhapsody was the best of all the tools at differentiating between pathogenic and non-pathogenic variants. Rhapsody incorporates dynamics information from elastic network models of protein structure, as well as PolyPhen-2 and EVMutation (conservation) scores using a machine learning approach. This broad range of sequence-, structure- and dynamics-based information could account for its success since the impact of a variant can arise from a multitude of different physio-chemical effects. Despite producing the best separation (Rhapsody) and describing features (SDM), structure-based tools are limited by the availability of the protein’s structure or a homologous structure. Although homology models are not always accurate for atomic level information, coarser detail such as whether residues are buried is likely to be reliable. Similarly, although Consurf can be used without a pdb structure it is dependent on homologous proteins having a structure. In this study we found using Consurf without a pdb structure was sufficient to predict pathogenicity based on whether a residue is likely to be buried or exposed, relating to its location in homologous proteins with known structures. Ultimately, these scores can only be used as a guide, and variant pathogenicity should be confirmed by segregation analysis and functional assays wherever possible, as rare non-pathogenic variants can also score highly.

Interrogation of gnomAD revealed a further three rare variants that were homozygous in at least one individual and could not be excluded as a cause of Oguchi disease based on allele frequency alone (Supplementary Table 3). We therefore calculated the same set of pathogenic prediction scores to assess the likely pathogenicity of these variants of unknown significance. The p.Met185Val variant, although rare in gnomAD, produced scores that were comparable with the non-pathogenic variants for each tool, suggesting it does not contribute to disease. However, both p.Ala353Ser and p.Ala387Val, produced scores that were compatible with pathogenicity, suggesting that the 2 individuals in gnomAD that are homozygous for these variants may have Oguchi disease. gnomAD contains exome and genome sequence data from over 130,000 unrelated individuals, with only those individuals affected by a severe paediatric disorder being removed. It is therefore possible that individuals with CSNB or Oguchi disease are present in the gnomAD database which could explain the presence of two homozygous, likely pathogenic variants.

In summary, we present seven novel variants in *GRK1* as a cause of Oguchi disease and perform an in-depth analysis of all *GRK1* variants in order to understand their contribution to disease. We describe three mechanisms likely to account for all *GRK1*-related disease, each of which ultimately result in a failure to phosphorylate light-activated rhodopsin. We assessed the ability of different *in silico* pathogenicity scores to differentiate pathogenic and non-pathogenic variants and found that Rhapsody out-performed all tools tested for this dataset. Finally, we have created an LOVD database into which all the known pathogenic variants and several common non-pathogenic variants described herein have been entered, in order to guide future variant interpretation in *GRK1*-associated disease.

## Accession Numbers

The variants reported in this paper have been submitted to the ClinVar database at the National Centre for Biotechnology Information (submission ref. SUB6701858).

## Supplemental Data

Supplemental Data includes two figures, three tables and the 3D homology model of Human GRK1 based upon the bovine GRK1 crystal structure (PDB: 4PNI).

## Acknowledgements

The authors would like to thank all the families for agreeing to participate in this study, and Roman A. Laskowski and Arun Prasad Pandurangan for assistance with VarSite and SDM respectively, and Ivet Bahar, Luca Ponzoni and the rest of the Bahar group for their help with Rhapsody. This study was supported by the European Retinal Disease Consortium (ERDC), the European Reference Network for Rare Eye Disease (ERN-EYE) and the Japan Eye Genetics Consortium (https://www.jegc.org/).

This study was funded by RP Fighting Blindness and Fight for Sight as part of the UK Inherited Retinal Disease Consortium (RP Genome Project GR586) and the UK National Health Service (NHS) Foundation Trust, the National Institute for Health Research Biomedical Research Centre (NIHR-BRC) at Moorfields Eye Hospital and the NIHR-BRC at Great Ormond Street Institute of Child Health, Moorfields Eye Charity and the Great Britain Sasakawa Foundation. MG is a PhD student on the Wellcome Trust 4-year PhD student in The Astbury Centre funded by The University of Leeds. GA is supported by a Fight For Sight (UK) Early Career Investigator Award. Sarah Harris acknowledges the EPSRC for their support (EP/S0306971/1). This research was made possible through access to the data and findings generated by the 100,000 Genomes Project managed by Genomics England Limited (a wholly owned company of the Department of Health and Social Care). The 100,000 Genomes Project is funded by the National Institute for Health Research and NHS England. The Wellcome Trust, Cancer Research UK and the Medical Research Council have also funded research infrastructure. The 100,000 Genomes Project uses data provided by patients and collected by the National Health Service as part of their care and support.

Members of the UK Inherited Retinal Disease Consortium include: Graeme C. Black, DPhil, FRCOphth (study chair); Georgina Hall, MSc; Stuart Ingram, BSc; Rachel L. Taylor, PhD; Panagiotis Sergouniotis, PhD; Andrew R. Webster, MD(Res), FRCOphth; Alison J. Hardcastle, PhD; Michel Michaelides, MD(Res), FRCOphth; Nikolas Pontikos, PhD; Michael Cheetham, PhD; Gavin Arno, PhD; Alessia Fiorentino, PhD; Chris F. Inglehearn, PhD; Carmel Toomes, PhD; Manir Ali, PhD; Martin McKibbin, FRCOphth; James A. Poulter, PhD; Kamron N. Khan, PhD, FRCOphth; Claire E.L. Smith, PhD; Susan Downes, MD, FRCOphth; Jing Yu, PhD and Veronica van Heyningen, PhD.

Members of the Genomics England Research Consortium include: Ambrose J. C., Arumugam P., Baple E. L., Bleda M., Boardman-Pretty F., Boissiere J. M., Boustred C. R., Brittain H., Caulfield M. J., Chan G. C., Craig C. E. H., Daugherty L. C., de Burca A., Devereau, A., Elgar G., Foulger R. E., Fowler T., Furió-Tarí P., Hackett J. M., Halai D., Hamblin A., Henderson S., Holman J. E., Hubbard T. J. P., Ibáñez K., Jackson R., Jones L. J., Kasperaviciute D., Kayikci M., Lahnstein L., Lawson K., Leigh S. E. A., Leong I. U. S., Lopez F. J., Maleady-Crowe F., Mason J., McDonagh E. M., Moutsianas L., Mueller M., Murugaesu N., Need A. C., Odhams C. A., Patch C., Perez-Gil D., Polychronopoulos D., Pullinger J., Rahim T., Rendon A., Riesgo-Ferreiro P., Rogers T., Ryten M., Savage K., Sawant K., Scott R. H., Siddiq A., Sieghart A., Smedley D., Smith K. R., Sosinsky A., Spooner W., Stevens H. E., Stuckey A., Sultana R., Thomas E. R. A., Thompson S. R., Tregidgo C., Tucci A., Walsh E., Watters, S. A., Welland M. J., Williams E., Witkowska K., Wood S. M., Zarowiecki M.

