## Supplementary Figure 1 for "New pathogenic variants and insights into pathogenic mechanisms in GRK1-related Oguchi disease"

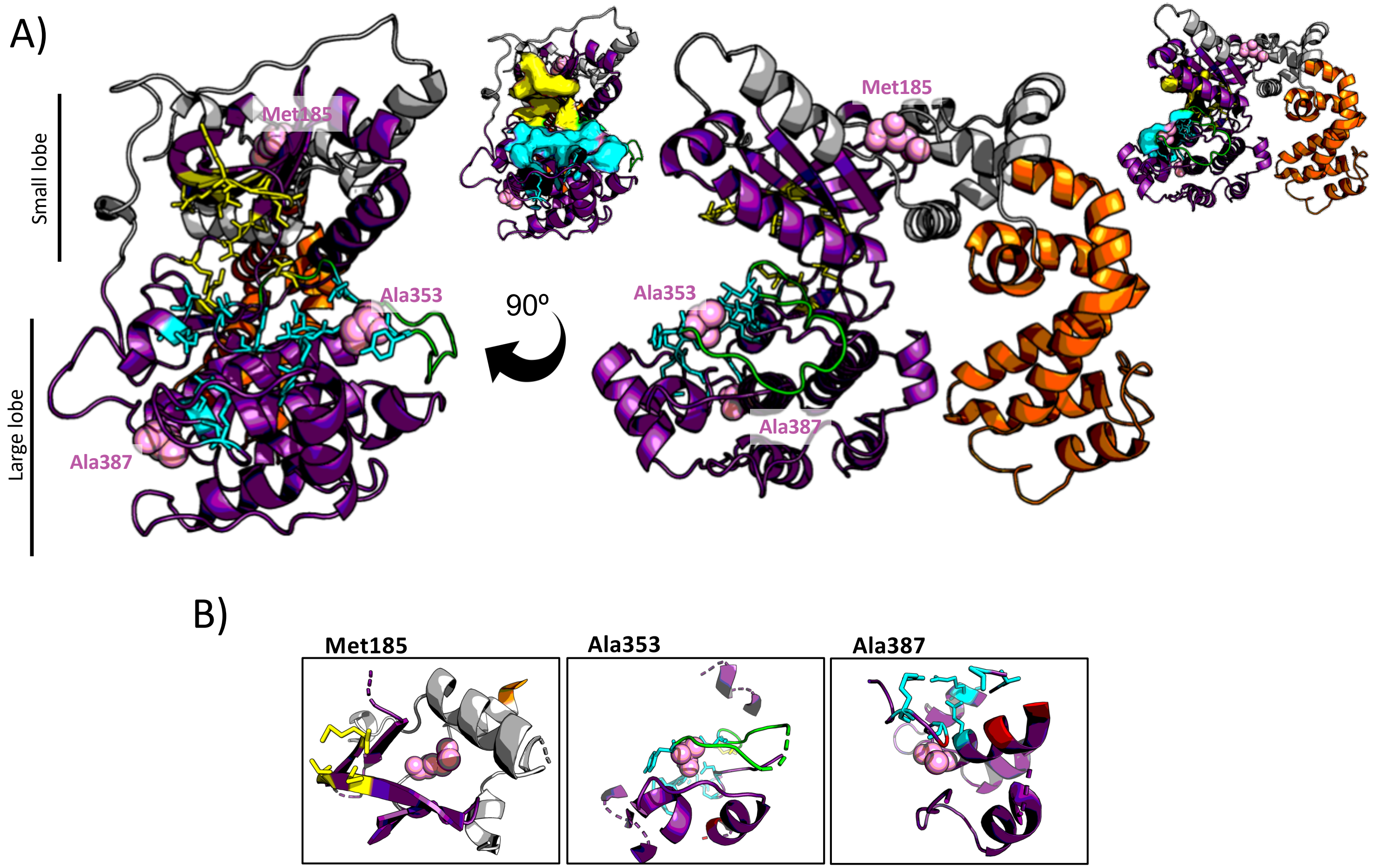


**Supplementary Figure 1** – Missense variants of unknown significance location in homology model of GRK1. Model of human GRK1 based upon bovine GRK1 structure (PDB: 4PNI). Wildtype residues of variants of unknown significance are labelled with pink spheres. The protein kinase domain is coloured in purple, regulator of G-protein signalling homology (RH) domain in orange, activation loop in green, ATP binding residues as yellow sticks and polypeptide substrate binding residues as cyan sticks. (A) Visualisation of variant location in overall structure. The smaller representations to the right, with the ATP and polypeptide binding surface displayed, demonstrate how intramolecular residue changes may deform the binding site surface. (B) A focus on residue location and local interactions, displaying only features within 15Å of the residue of interest.
