## Supplementary Figure 2 for "New pathogenic variants and insights into pathogenic mechanisms in GRK1-related Oguchi disease"

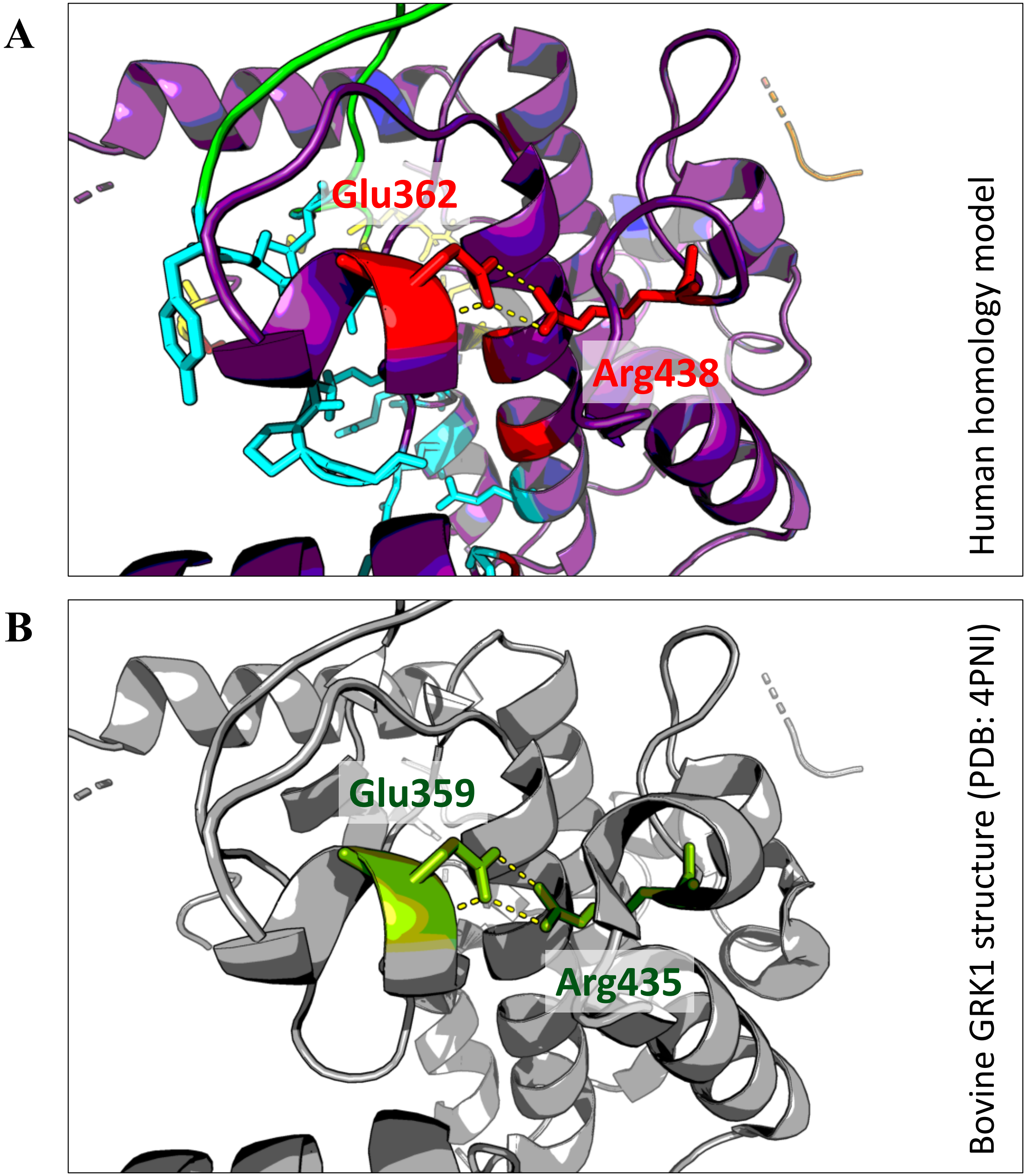


**Supplementary Figure 2 – Mutations at Glu362 and Arg438 are likely to affect GRK1 structure through disruption of a salt bridge interaction.**

A) Human homology model of GRK1 showing residues Glu362 and Arg438 interacting via a salt bridge. Substitution of either residue has been identified as a cause of Oguchi disease.

B) The homologous region in the bovine GRK1 crystal structure, showing the same residues (Glu359 and Arg435) interacting via a salt bridge.
