## Supplementary Data for "New pathogenic variants and insights into pathogenic mechanisms in GRK1-related Oguchi disease"

**Novel mutations and in-depth structural analyses provide new insights into pathogenic mechanisms underlying GRK1-related Oguchi syndrome**

James A. Poulter^1,^**^*^**, Molly S. C. Gravett^2,*^, Rachel L. Taylor^3^, Kaoru Fujinami^4-7^, Julie De Zaeytijd^8^, James Bellingham^6^, Atta Ur Rehman^9^, Takaaki Hayashi^10^, Mineo Kondo^11^, Abdur Rehman^12^, Muhammad Ansar^13^, Dan Donnelly^14^, Carmel Toomes^1^, Manir Ali^1^, UK Inherited Retinal Disease Consortium, Genomics England Research Consortium, Elfride De Baere^8^, Bart P. Leroy^8,15^, Nigel P. Davies^16^, Robert H. Henderson^17^, Andrew R. Webster^5,6^, Carlo Rivolta^18-20^, Omar A. Mahroo^5,6^, Gavin Arno^4-6^, Graeme C. Black^3,21^, Martin McKibbin^1,22^, Sarah A. Harris^23,**^, Kamron N. Khan^1,22,**^ and Chris F. Inglehearn^1,**^.

^1^ Division of Molecular Medicine, Leeds Institute of Medical Research, University of Leeds, Leeds, UK. ^2^ School of Molecular and Cellular Biology, University of Leeds, UK. ^3^ Division of Evolution and Genomic Sciences, School of Biological Sciences, Faculty of Biology, Medicines and Health, University of Manchester, Manchester, UK. ^4^ National Institute of Sensory Organs, National Hospital Organization Tokyo Medical Centre, Tokyo, Japan.

^5^ Moorfields Eye Hospital, London, UK. ^6^ UCL Institute of Ophthalmology, London, UK.

^7^ Keio University School of Medicine, Tokyo, Japan. ^8^ Ghent University, Ghent, Belgium.

^9^ Division of Genetic Medicine, Centre Hospitalier Universitaire Vaudois (CHUV) University of Lausanne, Switzerland. ^10^ The Jikei University School of Medicine, Tokyo, Japan. ^11^ Mie University Graduate School of Medicine, Mie, Japan. ^12^ Department of Genetics, Faculty of Science, Hazara University Mansehra, Pakistan. ^13^ Clinical Research Center, Institute of Molecular and Clinical Ophthalmology Basel (IOB), Basel, Switzerland. ^14^ School of Biology, University of Leeds, UK. ^15^ Children’s Hospital of Philadelphia, Philadelphia, PA, USA. ^16^ St Thomas’s Hospital, London, UK. ^17^ Department of Ophthalmology, Great Ormond Street Hospital, London, UK. ^18^ Department of Genetics and Genome Biology, University of Leicester, Leicester, United Kingdom. ^19^ Clinical Research Center, Institute of Molecular and Clinical Ophthalmology Basel (IOB), Basel, Switzerland. ^20^ Department of Ophthalmology, University Hospital Basel, Switzerland. ^21^ Manchester Centre for Genomic Medicine, Saint Mary’s Hospital, Manchester University NHS Foundation Trust, Manchester, UK. ^22^ Leeds Teaching Hospitals NHS Trust, St James’ University Hospital, Leeds, UK.

^23^ School of Physics, University of Leeds, UK. ^*^ These authors contributed equally to this work. ^**^These authors contributed equally to this work.

**Supplementary Data - Contents**

**Supplementary Table 1.** GRK1 variants with at least 1 homozygous individual present in gnomAD but considered too common to cause Oguchi disease.

**Supplementary Table 2.** Variant bioinformatic scores and residue property changes for individual missense variants.

**Supplementary Table 3.** Bioinformatic and structural scoring of rare homozygous variants of unknown clinical significance in gnomAD.

| **cDNA variant** | **Exon** | **Protein variant** | **rs number** | **MAF** | **Domain** | **No. of Hom** | **PolyPhen-2** | **CADD (v1.3)** |
| --- | --- | --- | --- | --- | --- | --- | --- | --- |
| c.160C>T | 1 | p.Leu54Phe | rs200774682 | 367/280268  (1.31e-3) | RH domain | 2^b^ | 0.273  (Benign) | 13.52 |
| c.290C>T | 1 | p.Thr97Met | rs116680839 | 490/280500  (1.75e-3) | RH domain | 2^a^ | 0.065  (Benign) | 14.76 |
| c.722T>C | 2 | p.Ile241Thr | rs116539667 | 841/280698  (3.00e-3) | Kinase | 11^a^ | 0.992  (Prob. D) | 24.7 |
| c.1390G>C | 6 | p.Glu464Gln | rs559629294 | 79/172662  (4.58e-4) | Kinase | 1^a^ | 0.619  (Poss. D) | 26.3 |
| c.1607C>T | 7 | p.Ser536Leu | rs553969577 | 96/172916  (5.55e-4) | C-terminus | 1^b^ | 0.047  (Benign) | 11.8 |

**Supplementary Table 1. GRK1 variants with at least 1 homozygous individual present in gnomAD but considered too common to cause Oguchi disease.** Genome position is based on the GRCh37 Human Genome, with variants annotated to NM_002929 (cDNA) and NP_002920 (protein). Values for minor allele frequency (MAF) and number of homozygotes (Hom) were obtained from gnomAD v2.1.1 [updated 8^th^ Jan 2020]. CADD scored are based on CADD v1.3. **^a^** All homozygotes originate from Africa. **^b^** All homozygotes originate from South Asia.

|  | **Protein variant** | **WT SSE** | **WT residue depth** | **WT OSP** | **Mw change** | **Change in pI** | **Hydrophobicity** | **Rhapsody** | **Consurf normalised score** | **Consurf buried/**  **exposed** | **Consurf predicted role** |
| --- | --- | --- | --- | --- | --- | --- | --- | --- | --- | --- | --- |
| Pathogenic variants | p.Leu157Pro | H | 8.1 | 0.61 | 16.04 | 0.32 | 0.98 | 0.86 | -0.985 | b | s |
|  | p.Gly199Arg | E | 3.5 | 0.3 | 99.14 | 4.79 | 1.01 | 0.76 | -0.915 | e | f |
|  | p.Leu308Pro | H | 10 | 0.55 | 16.04 | 0.32 | 0.98 | 0.93 | -0.940 | b | s |
|  | p.Glu362Lys | H | 5.5 | 0.5 | 0.94 | 6.52 | 0.35 | 0.72 | -1.112 | e | f |
|  | p.Ala377Pro | H | 11.8 | 0.54 | 26.04 | 0.30 | 0.41 | 0.69 | -0.539 | b | - |
|  | p.Val380Phe | H | 9.9 | 0.57 | 14.98 | 3.19 | 1.99 | 0.83 | -0.884 | b | s |
|  | p.Val380Asp | H | 9.9 | 0.57 | 48.06 | 0.48 | 0.57 | 0.75 | -0.884 | b | s |
|  | p.Pro391His | - | 7.7 | 0.51 | 40.02 | 1.29 | 0.59 | 0.82 | -0.891 | e | f |
|  | p.Arg438Cys | - | 7.5 | 0.55 | 53.04 | 5.69 | 2.55 | 0.75 | -1.022 | e | f |
| Non-pathogenic variants | p.Leu54Phe | - | 4.4 | 0.32 | 34.02 | 0.50 | 0.09 | 0.34 | 0.968 | b | - |
|  | p.Thr97Met | H | 4 | 0.39 | 30.09 | 0.14 | 0.97 | 0.17 | 0.899 | e | - |
|  | p.Ile241Thr | H | 6.2 | 0.48 | 12.06 | 0.42 | 1.54 | 0.42 | -0.610 | b | - |
|  | p.Glu464Gln | H | 3.6 | 0.26 | 0.99 | 2.75 | 0.04 | 0.13 | -0.611 | e | - |
|  | p.Ser536Leu | - | 3 | 0.04 | 26.08 | 0.30 | 1.74 | 0.32 | 1.404 | e | - |

**Supplementary Table 2 – Variant bioinformatic scores and residue property changes for individual missense variants.** Wildtype (WT) secondary structure location, WT residue depth and occluded surface packing (OSP) scores taken from SDM, Mw change and change in pI, hydrophobicity change taken from VarSite, Rhapsody scores from the Rhapsody webserver, and Consurf scores and interpretation calculated using Consurf. WT = wildtype, SSE = secondary structure location, OSP = occluded surface packing, Mw = molecular weight, H = alpha helical, E = beta hairpin, b = buried, e = exposed, s = structural (highly conserved and buried), f = functional (highly conserved and exposed), **-** = no effect predicted.

| **DNA variant** | **Exon** | **Protein variant** | **rs number** | **gnomAD Frequency** | **Domain** | **PolyPhen-2** | **CADD (v1.3)** | **Rhapsody** | **WT SSE** | **WT residue depth** | **WT OSP** | **Consurf normalised score** | **Consurf buried/**  **exposed** | **Consurf predicted role** |
| --- | --- | --- | --- | --- | --- | --- | --- | --- | --- | --- | --- | --- | --- | --- |
| ***Pathogenic variants (range)*** | | | | | | ***0.986-1.00*** | ***24.7-34.0*** | ***0.69-0.93*** |  | ***3.5-11.8*** | ***0.3-0.57*** | ***-1.112-(-)0.539*** |  |  |
| ***Non-pathogenic variants (range)*** | | | | | | ***0.010-0.986*** | ***11.80-26.3*** | ***0.13-0.42*** |  | ***3.0-6.2*** | ***0.26-0.48*** | ***-0.611-1.404*** |  |  |
| c.553A>G | 1 | p.Met185Val | rs564727279 | 20/270,752 | Kinase | 0.001  (Benign) | 0.001 | 0.05  (Neutral) | - | 6.29 | 0.428 | -0.243 | e | - |
| c.1057G>T | 4 | p.Ala353Ser | N/A | 2/142,244 | Kinase | 1.00  (Prob. D) | 27.8 | 0.67  (Deleterious) | - | 3.92 | 0.328 | -0.962 | e | f |
| c.1160C>T | 5 | p.Ala387Val | rs766150912 | 30/172,640 | Kinase | 0.992  (Prob. D) | 34.0 | 0.67  (Deleterious) | H | 5.52 | 0.480 | -0.364 | e | - |

**Supplementary Table 3. Bioinformatic and structural scoring of rare homozygous variants of unknown clinical significance in gnomAD.** All mutations are shown relative to Human genome reference GRCh37, and GRK1 references NM_002929 and NP_002920. The precise start and end locations for each domain in GRK1 were obtained from Lodowski et al. (2006). Prob. D = Probably Damaging as scored by PolyPhen2. WT = wildtype, SSE = secondary structure location, OSP = occluded surface packing, H = alpha helical, b = buried, e = exposed, s = structural (highly conserved and buried), f = functional (highly conserved and exposed), **-** = no effect predicted. The range of scores for pathogenic and non-pathogenic variants are taken from Table 1 and Supplementary Table 2.
